# Toward a More Accurate 3D Atlas of *C. elegans* Neurons

**DOI:** 10.1101/2021.06.09.447813

**Authors:** Michael Skuhersky, Tailin Wu, Eviatar Yemini, Edward Boyden, Max Tegmark

## Abstract

Determining cell identity in volumetric images of tagged neuronal nuclei is an ongoing challenge in contemporary neuroscience. Frequently, cell identity is determined by aligning and matching tags to an “atlas” of labeled neuronal positions and other identifying characteristics. Previous analyses of such *C. elegans* datasets have been hampered by the limited accuracy of such atlases, especially for neurons present in the ventral nerve cord, and also by time-consuming manual elements of the alignment process. We present a novel automated alignment method for sparse and incomplete point clouds of the sort resulting from typical *C. elegans* fluorescence microscopy datasets. This method involves a tunable learning parameter and a kernel that enforces biologically realistic deformation. We also present a pipeline for creating alignment atlases from datasets of the recently developed NeuroPAL transgene. In combination, these advances allow us to label neurons in volumetric images with confidence much higher than previous methods. We release, to the best of our knowledge, the most complete *C. elegans* 3D positional neuron atlas, encapsulating positional variability derived from 7 animals, for the purposes of cell-type identity prediction for myriad applications (e.g., imaging neuronal activity, gene expression, and cell-fate).

## Introduction

The nematode *C. elegans* is among the most-studied animals in neuroscience, and remains the only multicellular organism with a fully mapped connectome. Capable of exhibiting complex behaviors despite having as few as 302 neurons, it has provided an abundance of neuroscientific insights. The goal of this paper is to further improve the scientific utility of *C. elegans* by enabling more accurate and automated identification of its neurons, that are either unlabeled, or have been labeled by one or more colors that encode information regarding identity.

### Limitations of the original *C.elegans* connectome

The electron micrograph (EM) reconstruction of the *C. elegans* nervous system and its connectome were first fully described in a seminal 1986 paper (***White et al., 1986***). This was an invaluable technical achievement requiring a decade of work to hand-trace every neuron and connection from the EM sections. Due to technical limitations in preparing worm samples, this nervous system reconstruction was derived from a mosaic of overlapping sections from five individual worms. These five worms consist of three adult hermaphrodites, a fourth larval stage (L4) animal, and one adult male. Thus, in combination, this reconstruction provides a generalized view of the worm’s nervous system.

While a generalized view of the worm’s nervous system has proven valuable to the field, it lacks representation for the idiosyncrasies found among individual worms. Moreover, preparations for EM imaging can introduce non-linear distortions. Preparing worms for EM imaging requires that they be first physically sliced open so that the fixative may bypass the impermeable cuticle (***Hall et al., 2002-2021, 2012***). As the worm interior is under a higher pressure, breaching the cuticle results in morphological changes, and the commonly used EM fixative osmium tetroxide has also been shown to alter morphology (***Zhang et al., 2017***). In the original reconstruction of the worm nervous system, animals were serially sectioned into approximately 50 nm thick slices. The combination of these steps yielded distortions that required correction when unifying worm sections into a general visual representation of its nervous system. As a result, substantial manual correction was introduced when generating this canonical nervous system illustration. These corrections and the multi-animal synthesis thus present a generalized view of the worm’s nervous system, albeit one that lacks quantification of neuron positions and measurements of their variability to recapitulate individual idiosyncrasies present within the population.

### Existing *C.elegans* atlases

This illustrative 1986 worm atlas has often been treated as canon, or at least as an atlas of sufficient quality to compare with contemporary data from modern and more reproducible measurement techniques. This is in large part due to it being the only atlas of its kind up until very recently (***Toyoshima et al., 2020; Yemini et al., 2021***). For example, Scholz *et al*. (***Scholz et al., 2018***) aligns this atlas to fluorescent imaging data to assign neuron identity. Unfortunately, using this generalized illustrative atlas for neural identification purposes can lead to unlabeled and even mislabeled neurons. This is largely due to the density of neurons in various ganglia and ambiguities in atlas matching that are present as a result. In addition to these limitations and those listed in the previous section, a 2004 version of this atlas, commonly used in papers that make use of *C. elegans* neuron positions, was produced by further processing the original version so as to translate the 1986 illustration into semi-quantifiable measurements. This version was produced by Choe *et al*. (***Choe et al., 2004***) and popularized by Kaiser *et al*. (***Kaiser and Hilgetag, 2006***). It tabulates neural positions that were measured by directly scanning and tracing the physical 1986 paper. The 2004 atlas consists of only 277 of the 302 neurons found in adult hermaphrodites. Crucially, due to the 2D geometry of the scanned and processed original paper, the 2004 atlas is also 2D. The third dimension in modern 3D datasets is therefore typically discarded before matching against this 2D atlas and may further increase the misidentification rate. In total, these limitations motivate the creation of a representative 3D atlas of *C. elegans* neuron positions and their variability.

In the decades since the original EM connectome was published (***White et al., 1986***), many individual *C. elegans* neurons have been further studied and characterized. Their updated information has been incorporated into the widely used *C. elegans Atlas* (***Hall et al., 2007***), and the contents of this reference text have been artistically assembled into the popular and influential OpenWorm project (***Szigeti et al., 2014; Gleeson et al., 2018***). OpenWorm presents information for individual neurons and their connectivity, assembled into a 3D approximation of an adult hermaphrodite worm. However, the inherent positional variability of neurons in the head and tail is now readily apparent from multicolor *C. elegans* strains, designed for neural identification, that were used to measure neuron positions and their variability across multiple animals (***Toyoshima et al., 2020; Varol et al., 2020; Yemini et al., 2021***). These strains and concurrent algorithmic advances demonstrate how fluorescent-protein barcodes can be used to accurately determine neuron identities in volumetric images (***Bubnis et al., 2019; Nejatbakhsh et al., 2020; Mena et al., 2020; Chaudhary et al., 2021; Yu et al., 2021; Wen et al., 2021***).

The strong interest in *C. elegans* research coupled with great recent progress in all aspects of *C. elegans* imaging makes it timely to develop improved tools for whole-nervous-system *C. elegans* neuron identification and make them available to the community. This is the goal of the present paper. The rest of this paper is organized as follows: we present our improved alignment method in Materials and Methods together with a pipeline for creating improved atlases of whole-worm neuron positions. We then compare the neuron identification accuracies of various methods in Results, applying them to simulated data and then to our new atlas. We then summarize these results in our Discussion.

**Figure 1.**
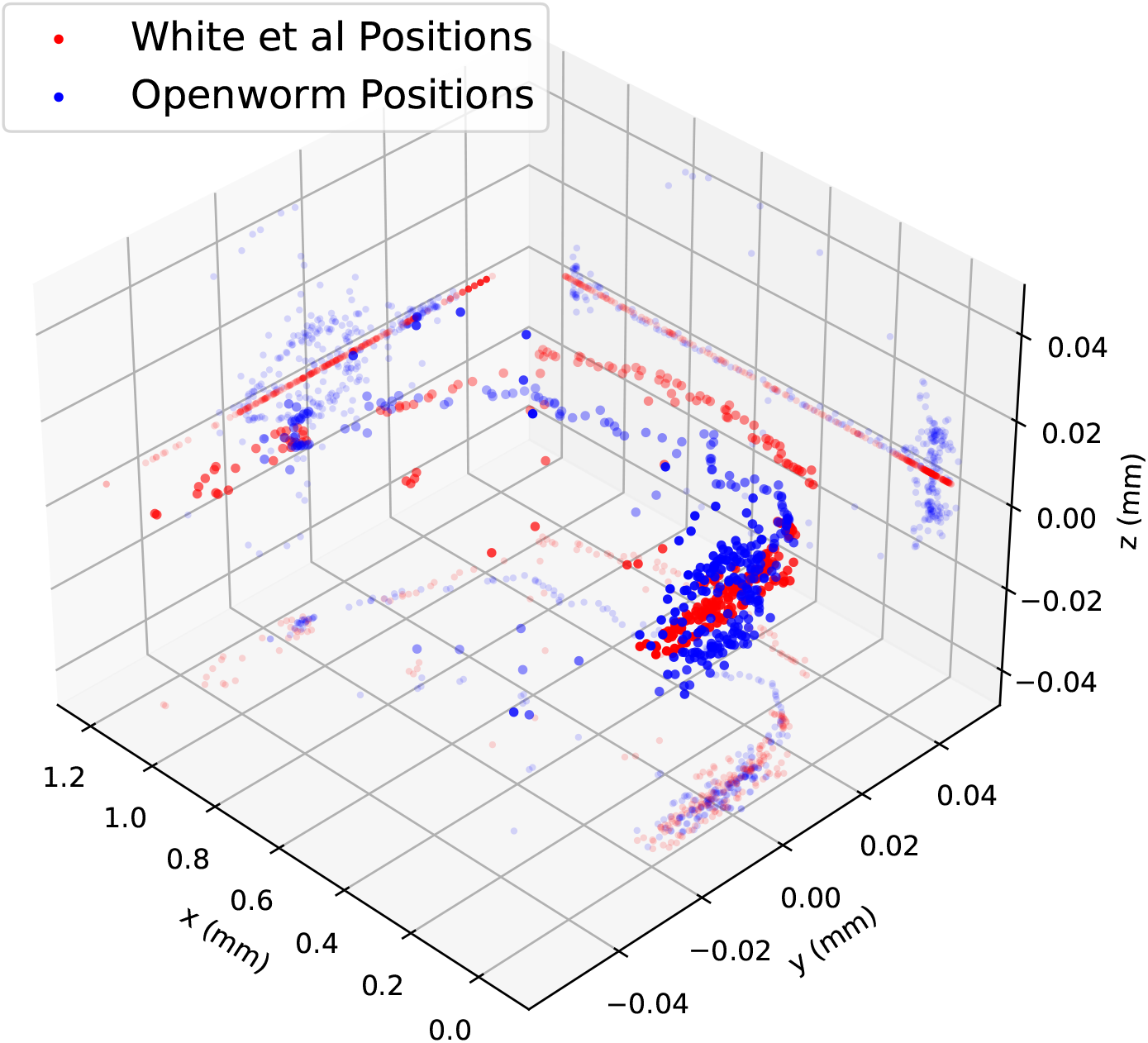
*C. elegans* atlases from White et al. (blue) and OpenWorm (red), after straightening and uniform scaling using a corrected version of (***Marblestone, 2016***), with axial projections, unequal scaling.

## Materials and Methods

### New NeuroPAL-derived Atlas

Our construction of the NeuroPAL atlas consists of three steps:

1. Preparing and imaging NeuroPAL worms and turning each image into a pair of measured quantities as in Preparing and imaging the NeuroPAL worms: a point cloud (a position vector and optional color information for each neuron) and a hull outline (a smooth closed 2D curve).
2. Canonicalizing each pair by normalizing the hull into a standard straightened hull, along with the point cloud in the associated canonical 3D space.
3. Combining point clouds from multiple worms into a single point cloud atlas.

Our constructed NeuroPAL atlas is shown in Atlas construction and presented as a coordinate table in the Appendix.

**Figure 2.**
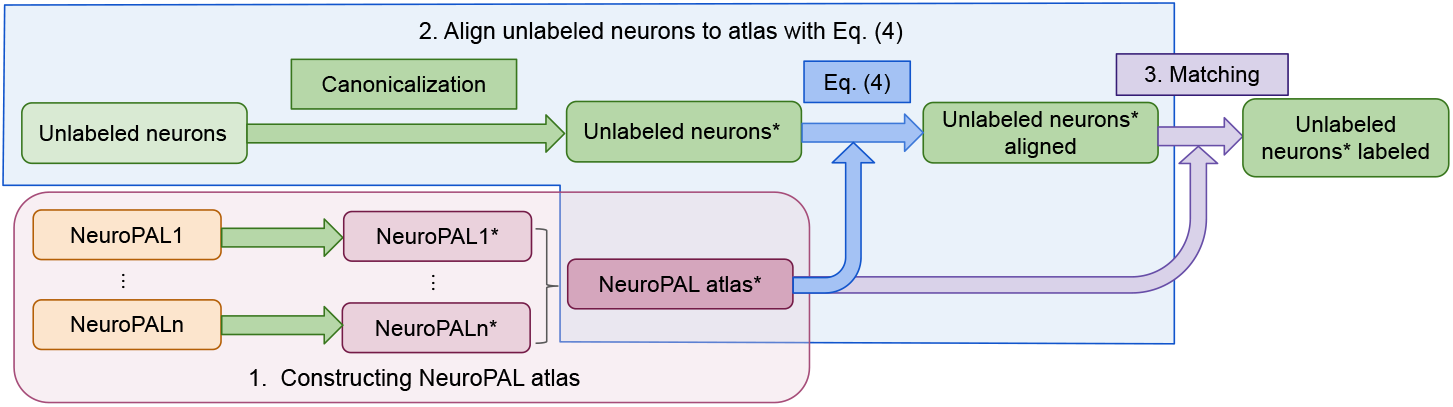
Diagram of our NeuroPAL atlas construction and alignment pipeline, with each algorithm labeled. The * denotes the neuron positions after canonicalization.

#### Preparing and imaging the NeuroPAL worms

Brainbow (***Weissman and Pan, 2015***) is a stochastic technique that has been used to differentiate individual neurons from neighboring ones by expressing unique ratios of red, green, and blue fluorescent proteins. Unfortunately, Brainbow coloring is generated randomly and thus the colors cannot be used to identify neuron types. In contrast, the recent NeuroPAL *C. elegans* transgene introduces an alternative deterministic technique that reveals the unique identity of each individual neuron, at all larval stages of both worm sexes (hermaphrodites and males) (***Yemini et al., 2021; Tekieli et al., 2021***). NeuroPAL worms have an identical and invariant colormap by way of the stereotyped expression of four distinguishable fluorophores. These four fluorophores leave the green channel free, and thus it can be used to map gene expression using GFP, CFP, or YFP based gene reporters, or to measure dynamic neural activity using GCaMP.

A complicating factor in position-derived identification is that individual neuron positions are known to be locally variable between different worms (***Wall et al., 2007; Toyoshima et al., 2020; Yemini et al., 2021***). This is caused partly by *C. elegans* size and shape differences, and partly because of inherent positional variance that may be so extreme as to have nearby neurons switch relative positions.

Adult *C. elegans* were imaged in accordance with the methods as described in (***Yemini et al., 2021***), and the images were processed with a semiautomatic method using the software developed by (***Yemini et al., 2021***). Neuron positions were detected by the software, and any required corrections were made manually. The software predicted neuron identities for the head and tail of each worm, and these were manually checked and relabeled as needed. Since there were no prior statistical atlases for neurons in the worm midbody, the identities of these neurons were manually annotated for each worm. The original 1986 nervous system reconstruction (***White et al., 1986***) was assembled using a mosaic of 5 overlapping image sections, each corresponding to part of the worm, that when pieced together would represent entire nervous system. Each section was taken from one of 5 worms representing a mixture of age and sex: 3 adult hermaphrodites, 1 L4, and 1 adult male. In our atlas, to maintain a generalized representation of the nervous system we used 7 worms: 1 adult hermaphrodite, 4 young-adult hermaphrodites, 1 L4 hermaphrodite, and 1 adult male. Here, each individual worm has neuron positional information represented for its whole body, rather than only a body section, thus contributing a holistic representation of the nervous system from each of these animals. All 7 worms were positioned on an agar pad for imaging such that their left-right axis extended between the glass coverslip and slide. As a result, the worm samples and representative left-right axis may have been slightly compressed between these two surfaces. Similarly, the dorsal-ventral and potentially the anterior-posterior axes may have been slightly elongated as a result of this compression.

#### Straightening method

For successful alignment, canonicalization is essential in ensuring that the neuron point clouds from different worms, that were imaged in different postures, orientations, and morphology, lie in the same canonical space to provide a reasonable starting point for neuron matching. Mathematically, canonicalization corresponds to a continuous and invertible 3D mapping that that gives all imaged worm hulls the same shape regardless of the proportions and bending state of the worms.

Several methods have been previously used for this problem of straightening worm hulls for eventual canonicalization (***Peng et al., 2008; Christensen et al., 2015***). Because worms are imaged under a coverslip and thus lie in a 2D plane, such methods customarily model the 3D mapping as a 2D mapping, leaving the vertical dimension unchanged. We make the same simplification, because our current data does not provide sufficiently accurate 3D hull determination (worm edges are too blurred in less-focused horizontal slices far above or below the midplane).

Most previous methods do not attempt to preserve volume, and pose challenges related to distorted straightening at the head and tail. This is unfortunate, since these dense areas, presumably responsible for a majority of the worm’s information processing, are the most important to get right for scientifically-relevant neuron identification. For example, we originally tested a simple 2D canonicalization where the new *x*-coordinate was defined as the distance along the worm midline estimated by skeletonizing the hull image, and the new y-coordinate was defined as the perpendicular distance to this midline. This scheme unfortunately resulted in problematic volume distortions associated with the contorted morphology of an unrestrained worm, and produced relatively poor neuron matching accuracy.

We therefore propose a worm canonicalization method which produces more consistent and biologically realistic point clouds. As explained below, the 2D mapping can be geometrically interpreted by filling the worm with inscribed circles as illustrated in Figure 3. We first make a hull map for each worm, by binarizing the slice from the *C. elegans z*-stack that represents the largest-area 2D hull of the worm (Figure 3, middle panel).

**Figure 3.**
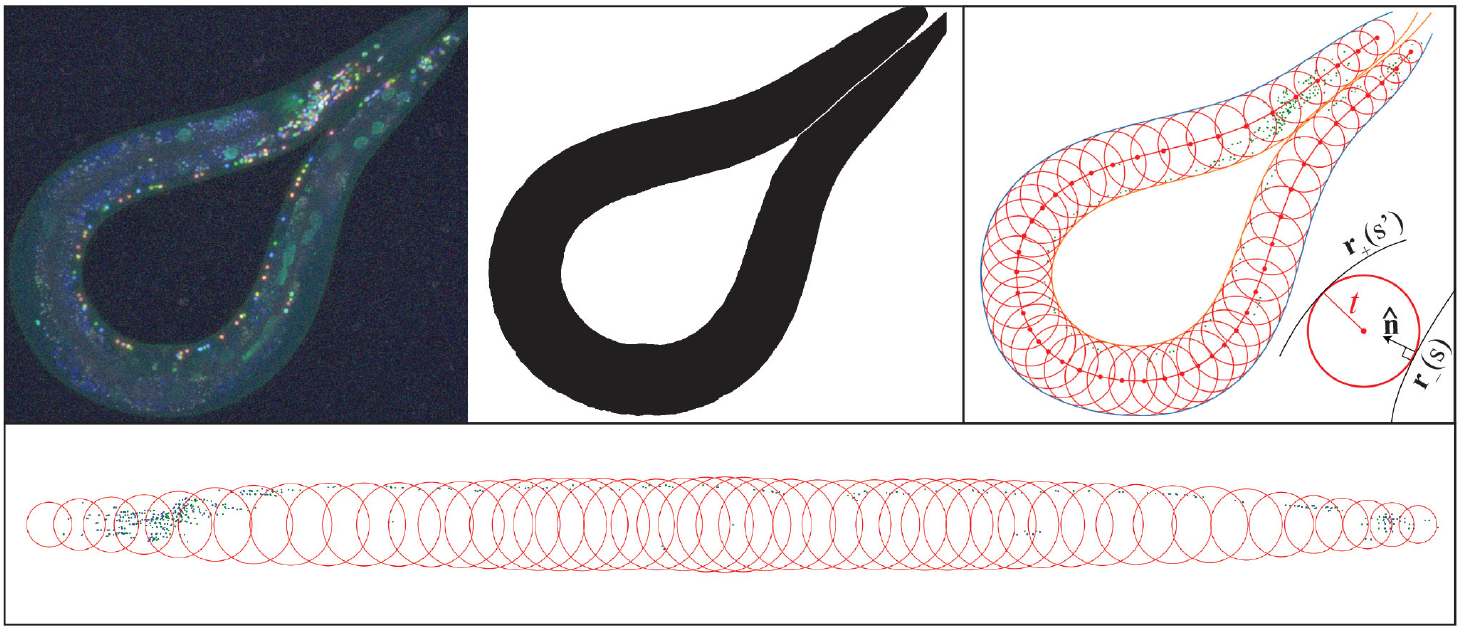
Starting with a worm image (top left), we first determine the hull (top center). Our algorithm then inscribes 1,000 tangent circles (top right) in the hull (for clarity, only 50 circles are shown here). Finally, the circle sequence is straightened, which defines the canonicalization mapping of their inscribed neuron positions (bottom).

We then split the worm hull boundary into two parameterized curves representing opposite edges, **r**_−_(*s*) and **r**_+_(*s*′), where *s* ∈ [0,1] and **r**_−_(*s*) runs clockwise from head to tail as *s* increases. We define these curves by cubic spline fits to the binarized hull image. As illustrated in Figure 3 (right panel), the distance from the point **r**_+_(*s*′) on one edge of the worm to a circle of radius *t* tangent to **r**_−_(*s*) on the other edge is then given by

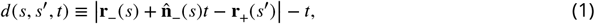

where 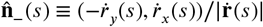 is the unit inward tangent vector at **r**_−_(*s*), the vector 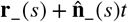 is the center of the aforementioned circle, and dots denote derivatives with respect to *s*. By numerically minimizing over *s*′, we find the distance *d*(*s, t*) from the circle to the opposite worm edge:

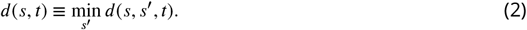

We numerically solve the equation *d*(*s, t*) = 0 to determine the radius *t*(*s*) where the circle is tangent to both worm edges. The variable *s* ∈ [0,1] thus parametrizes a continuous family of circles of radius *t*(*s*) centered at

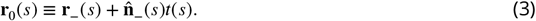

Finally, these circles are used to remap the *C. elegans* neurons using the methods illustrated in Fig. 3. For each worm, 1,000 equally spaced circles are inscribed. All worm-inscribed circles are then translated so that their midpoints lie on the *x*-axis, retaining the distances between adjacent circle midpoints, and rotated so that the worm midline corresponds to the x-axis (Fig. 3, bottom). Each circle thus defines an affine transformation into the canonical space. Each neuron is mapped into this canonical space by applying the affine transformation for each of the circles that inscribe it, and averaging the result. Finally, the straightened group of neurons are isotropically rescaled so as to occupy *x* ∈ [0*μm*, 800*μm*].

#### Atlas construction

After applying our canonicalization procedure to each individual worm and obtaining corresponding 3D neuron point clouds for each one, we combine these point clouds into a single NeuroPAL atlas by using the median position coordinates for each neuron. We use the median rather than the mean since it is more robust to outliers and resistant to overall shrinking. The resulting atlas is illustrated in Fig. 4, and Appendix 2 lists the 300 canonicalized neuron position coordinates. We do not provide data for the two CAN cells in our dataset, as previous broad investigations of panneuronal markers found none that solely express in neurons and also express in CAN (***Stefanakis et al., 2015***). All panneuronal markers that expressed in CAN also expressed in non-neuronal tissues such as epithelium, intestine, glands, or muscle. Non-neuronal tissues are often larger than neurons and fluorescent expression within them can can occlude neuronal imaging. Thus, to date, all whole-brain activity imaging as well as neural identification strains have used panneuronal markers that exclude CAN and, accordingly, these two cells are not represented in our dataset.

**Figure 4.**
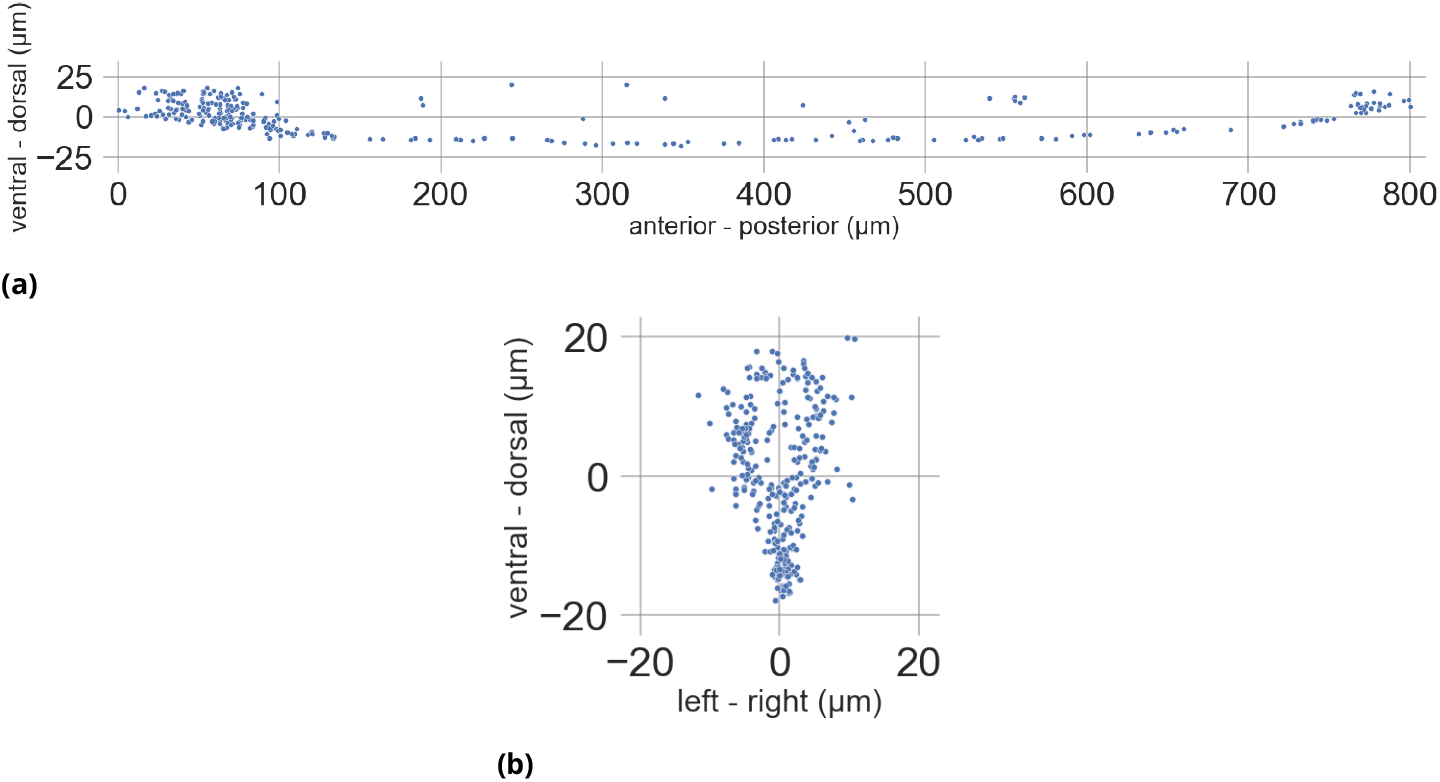
Sagittal (a) and transverse (b) views of our NeuroPal-derived atlas

### New Alignment Technique

We now turn to the challenge of aligning the neuron point cloud from an observed worm to a known atlas, so as to determine the likely identity of each observed neuron.

#### Existing Coherent Point Drift Alignment

Coherent Point Drift (***Myronenko and Song, 2010***) (CPD) is a probabilistic alignment method for sparse point clouds that has been extensively used to align *C. elegans* neuron positions (***Scholz et al., 2018; Toyoshima et al., 2020; Yu et al., 2021***). CPD represents the first point set by Gaussian mixture model (GMM) centroids and aligns to a second point set by maximizing the likelihood while forcing the points to move coherently as a group. In the non-rigid case, this is implemented as a motion over time guided by a velocity field determined by maximizing the GMM likelihood penalized by motion incoherence. CPD performs well on our *C. elegans* problem when the two point clouds undergo rigid-body rotation, translation and scaling, but as we will show below, has limited robustness in cases of cropping, size imbalance, and realistic biological deformation.

#### Colors

Since NeuroPAL is a fluorescent-labeling transgene that improves neuron distinguishability using several colors, it is highly desirable to exploit this color information to improve neuron matching. This begs the question: how many colors are needed for accurate alignment? With better alignment methods, are fewer colors sufficient? To quantify the relationship between color and accuracy, we report below a series of simulation results where each neuron is randomly assigned a simulated color.

#### Novel Generalized-mean Alignment

To improve the robustness of the alignment and address the issues in CPD, we introduce a novel *Generalized-Mean* (GM) alignment algorithm. For labeled neuron positions 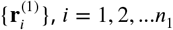 from reference worm 1 and unlabeled neuron positions 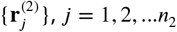 from worm 2 (*n_i_* denotes the number of neurons in the *i*^th^ worm), and optionally provided colors 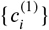 and 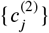, we introduce the following loss function to minimize:

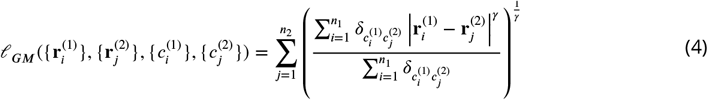

where 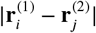 is the Euclidean distance between 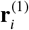 and 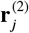, and the Kronecker delta 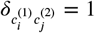 if the two colors are equal, vanishing otherwise. The generalized mean is defined by the hyperparameter *γ* < 0 which, as explained in (***Wu and Tegmark, 2019***), encourages pairing. More specifically, for each unlabeled neuron positions 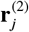, we have a generalized-mean of its distance to all the labeled neuron positions 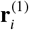 that have the same color (as enforced by 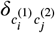 When *γ* < 0, the smaller the distance, the larger 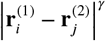 is. Furthermore, as proved in (***Wu and Tegmark, 2019***), this generalized-mean loss has the property that the smaller 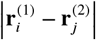 is among all the 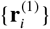, the larger the gradient is to drive them closer. Here *γ* tunes how much the algorithm is focusing on the smaller distances. For example, when *γ* → −∞, Eq. (4) reduces to 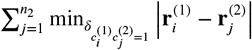 where each unlabeled 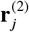 only focuses on the nearest labeled 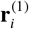 which can easily fall into a local minimum. However, allowing a *finite* negative *γ* also allows consideration of other potential parings that are not as near. In this paper, we set *γ* = −6, which we experimentally found to achieve a good balance between the aforementioned pairing effect and the desire for error correction whereby neurons at slightly larger distance produce enough of a gradient to be pushed together, which is crucial for avoiding getting trapped in suboptimal local minimal during the initial stages of training. In contrast, *γ* = 2 would correspond to *ℓ* being minus two times the log-likelihood of a Gaussian mixture model, making the loss similar to that of CPD with a Gaussian kernel.

### Algorithm testing framework

To identify unknown neurons, our pipeline proceeds as illustrated in Fig. 2. It first canonicalizes the unlabeled neurons, then aligns them with the canonicalized atlas as described above. The alignment function takes as input an unlabeled point cloud of neuron positions, an atlas point cloud with known neuron IDs (with optional neuron colors), and returns transformed positions for the unlabeled point cloud (step 2). Finally, in step 3, each neuron with transformed position is assigned an ID by the following procedure: for the *n*_2_ × *n*_1_ matrix of pairwise Euclidean distances between the *n*_2_ neurons in the unlabeled point cloud and the *n*_1_ neurons in the atlas, find the smallest element in the matrix, assign the corresponding ID, then delete this row and column in the matrix, and repeat. If colors are provided, each unlabeled neuron can only be assigned to a neuron in the atlas with the same color, so we choose the smallest element with unlabeled colors at each iteration.

We quantify the accuracy of all algorithms by testing them on simulated data where the ground truth is known. For these simulations, we use the OpenWorm dataset as the ground truth point cloud, distort it with simulated noise and biological deformations as described below, and finally measure which algorithms provide the most accurately reconstructed neuron identifications.

### Parameterizing more realistic worm deformation

Working in the above-defined canonical space, we express the relation between the neuron positions 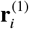 in reference worm 1 (our atlas, say) and observed neuron positions 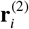 for a worm 2 as

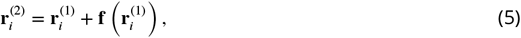

for a *deformation function* **f**(*r*) that may include a random component. Avery easy-to-simulate type of deformation is to simply add independent Gaussian noise to all coordinates of all neurons. This corresponds to treating all neuron positions 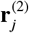 as independent parameters, and is tantamount to ignoring all biological constraints on tissue stretching.

We wish to regularize the problem to limit our analysis to more biologically plausible deformations, reflecting the known fact that positional variation between organisms exhibits correlation, whereby the deformation vector **f** is typically similar in direction and magnitude for adjacent neurons. In other words, we wish the deformation function **f**(**r**) to be relatively smooth, corresponding to nearby parts of the worm mainly shifting together as a coherent unit. We model the deformation field as a Gaussian mixture:

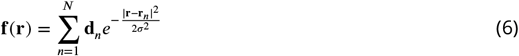

Here the deformation function is parametrized by a vector **p** of 6*N* + 1 parameters: the number *σ*, the components of the displacement vectors **d**_*n*_, and the displacement centers **r**_*n*_. A sample deformation field is visualized in Fig. 5. For better numerical stability, we add three redundant parameters in the form of an overall global displacement vector **s** added to **r**_*n*_.

**Figure 5.**
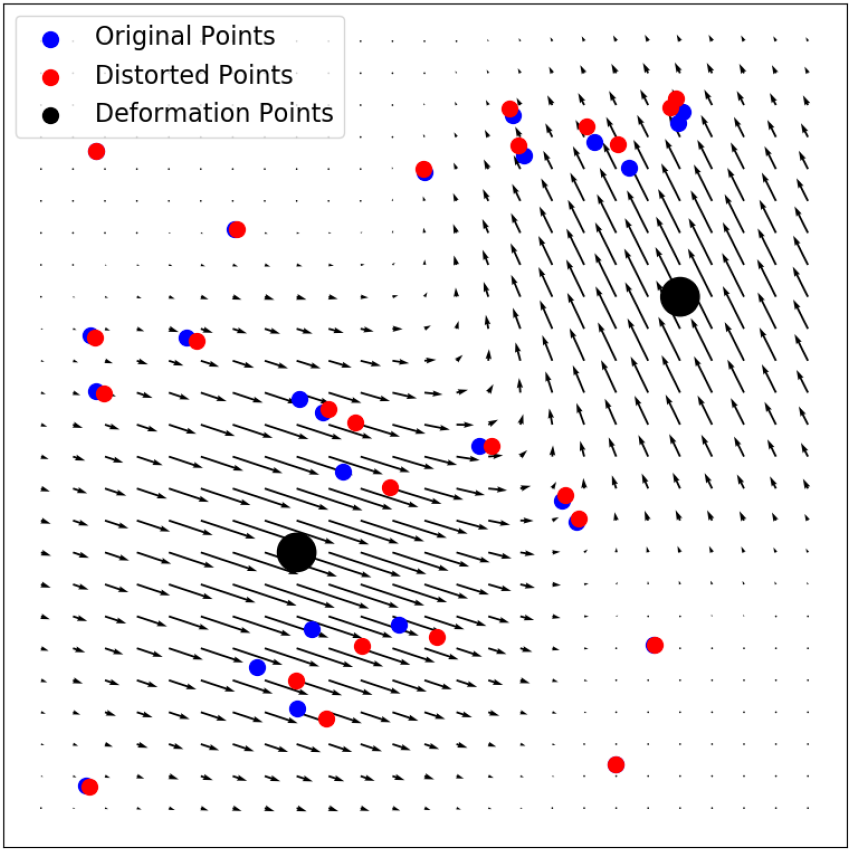
Illustration of simulated biological noise simulated with the “bio-realistic” deformation field.

### Hyperparameter Tuning

The performance of all tested alignment algorithms is highly hyperparameter dependent. To make a robust comparison of methods, we must ensure that for each algorithm we are using the optimal parameters for our use case. Therefore, we submit each algorithm to parallelized hyperparameter optimization.

Using the “Random Search” followed by the “Local Search” algorithms in the Sherpa (***Burke et al., 2020***) parameter tuning package, input parameters were tuned until alignment accuracy did not improve further. Here, alignment accuracy was defined as the average of performance per method in the tests Robustness to Cropping, Robustness to Dropout, and Robustness to Biological Noise. The determined optimal parameters (Appendix 1) for each algorithm were used in all subsequent testing.

## Results

We now compare the performance of our atlas and alignment methods with existing techniques. We designed experiments to answer the following questions: Firstly, how do our alignment method compare with existing methods against various types of adverse point cloud distortions? This is answered in Alignment performance on simulated data. Secondly, how does the new atlas perform compared to existing atlases? We answer this question in Testing alignment methods - Real Data by comparing the accuracy of aligning unlabeled neurons to existing and new atlases. Thirdly, since NeuroPAL and other labelling schemes use multiple colors to reveal identification, how many colors are necessary to achieve acceptable accuracy? In Testing alignment methods - Real Data, we also examine the effects of multiple colors on identification accuracy.

The following tests were devised: the point cloud positions of the 302 Openworm neurons were taken as a test set, as it can be assumed that these positions are reasonably representative of the morphology of actual *C. elegans* neurons (***Gleeson et al., 2018***). Then, a copy of this point cloud was made, and selected perturbations, designed to simulate real world experimental conditions were performed. Finally, the group of points was randomly perturbed by a single vector drawn from a uniform distibution between ±5 microns in each of the *x*, *y*, and *z* directions. The alignment algorithms were then used to align the perturbed neuron point cloud to the original one. We test alignment with GM-realistic (GM with Gaussian deformation centers as in defined in Eq. (6)), CPD-rigid (CPD with rigid transformation, as defined in Fig. 2 of (***Myronenko and Song, 2010***)), and CPD-deformable (CPD with deformable transformation, as defined in Fig. 4 of (***Myronenko and Song, 2010***)).

This procedure was repeated 40 times at each perturbation setting, with resulting accuracies averaged together to provide a final metric of accuracy.

### Alignment performance on simulated data

To simulate point clouds data similar in structure to those from real *C. elegans* images, we start with the OpenWorm atlas (***Gleeson et al., 2018***) and generate a point cloud of neuron positions with the ground-truth neuron IDs hidden from the alignment methods. A good alignment and identification pipeline should be robust to the above scenarios; therefore we sought to test the robustness of our alignment algorithm when deforming the Openworm point cloud dataset in these ways. In the following plots, results for optimally tuned parameters are shown.

#### Robustness to Cropping

Commonly, *C. elegans* activity research focuses on the head of the worm, due to microscope FOV or resolution reasons, or hypotheses aimed at the nerve ring. During alignment, the head neurons are usually matched to a larger atlas. Therefore, a good identification method must perform well when aligning a dataset of unlabeled neurons that may reference a cropped subset of the comparison atlas.

For the first test, we start with a full Openworm point cloud that has been isotropically rescaled so as to occupy *x* ∈ [−600*μm*, 200*μm*]. We choose all head neurons (located at *x* > 0*μm*) as the unlabeled neuron set, and also create copies of the Openworm point cloud with various *x* cropping thresholds to act as the comparison atlas. We perturb the larger atlas as a rigid group away from the head neurons by a single vector drawn from a uniform distibution between ±5 microns in each of the *x*, *y*, and *z* directions. Then we use the alignment algorithms to align the unlabeled head neurons to the various croppings of the larger atlas, to test how different algorithms are robust to the imbalance of the two neuron sets.

Fig. 6 shows the resulting alignment accuracy for GM-realistic, CPD-rigid, and CPD-deformable. We can see that the GM-realistic algorithm performs perfectly across all levels of imbalance, while the CPD methods, accuracy drops quickly when the atlas extends beyond −100*μm*. The robustness of our GM method comes from the fact that the loss in Eq. (4) has the soft “clamping” effect that is able to focus on well-matched pairs of neurons while ignoring pairs that are matched poorly. The CPD methods, in contrast, are more sensitive to those badly-matched pairs, so the performance quickly drops when the two neuron sets are imbalanced. We also became aware of differences in alignment performance when aligning two sets of head neurons, from aligning two sets of wholebody neurons. The whole-body neurons take up space that is elongated along one axis, whereas head neurons occupy a relatively tight spherical volume. Because of this, head-head alignments were found to be more likely to fail by getting stuck in false local minima with the CPD methods.

**Figure 6.**
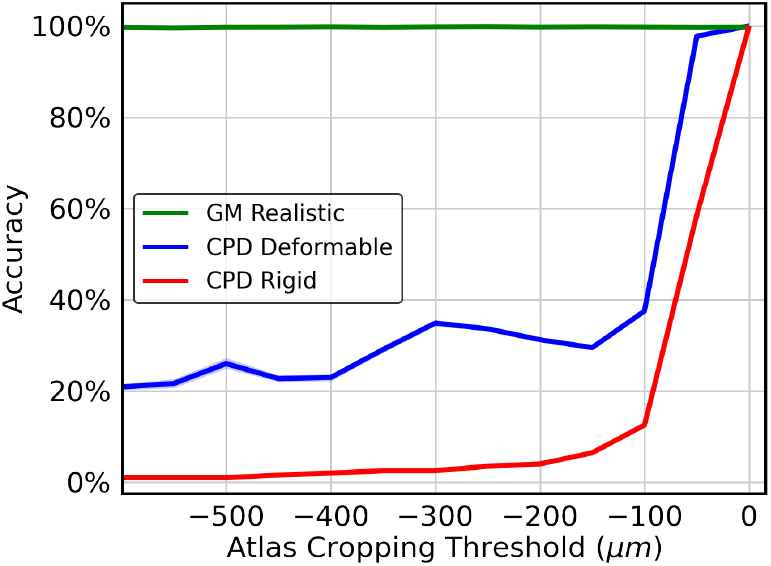
Cropping Performance (40 run average): Alignment accuracy after aligning OpenWorm data cropped to the head to OpenWorm data cropped at various *x*-coordinates and perturbed as a group in a random direction in *x*, *y*, and *z* each at up to 5 microns. At *x* = 0, the atlas cropping isolates the head.

**Figure 7.**
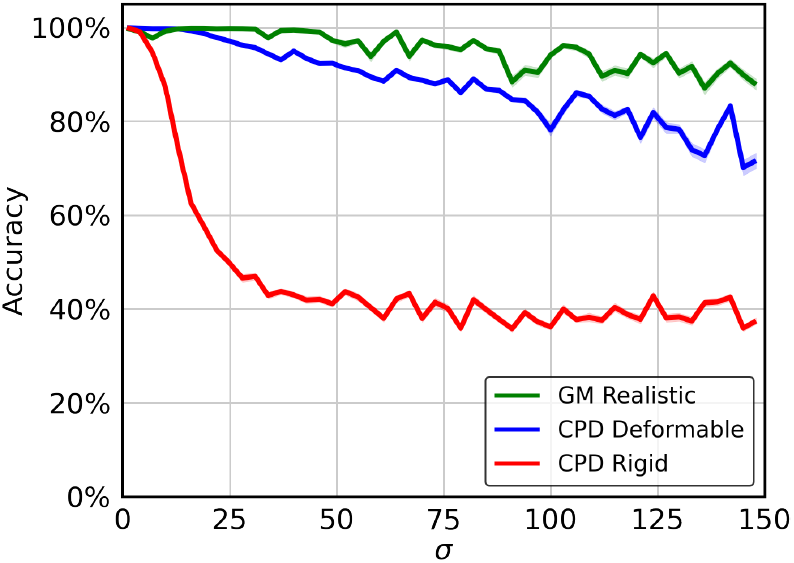
Biological Noise Performance (40 run average): Alignment accuracy after aligning OpenWorm data to OpenWorm data that has been deformed with our biological noise method at various influence radius *σ* and perturbed as a group in a random direction in x, y, and z each at up to 5 microns.

#### Robustness to Dropout

Fluorescent-protein expression that is used to identify neurons can at times be too dim to resolve. As a result of their dimness, these neurons cannot be detected or definitively identified. Such situations can occur due to many reasons, for example fast volumetric imaging methods that necessitate a tradeoff between speed and imaging quality. More generally, dimness is often found due to optical anisotropy present in various volumetric imaging techniques. A good identification method must therefore be robust to various levels of neural dropout.

For this, we start with a full OpenWorm point cloud. Then, we randomly remove a set number of neurons, and set the result as our unlabeled point cloud, with another copy of the OpenWorm point cloud as the comparison atlas. We perturb the larger atlas as a rigid group away from the head neurons by a single vector drawn from a uniform distibution between ±5 microns in each of the *x*, *y*, and *z* directions. Then we use the alignment algorithms to align the unlabeled set to the complete atlas.

Fig. 8 shows the resulting alignment accuracy for GM-realistic, CPD-rigid, and CPD-deformable. We can see that the GM-realistic algorithm performs the best in all cases of neuron removal, plateauing to 100% accuracy with only 80 out of 300 neurons remaining. The CPD methods are more prone to misidentification by getting stuck in suboptimal local minima.

**Figure 8.**
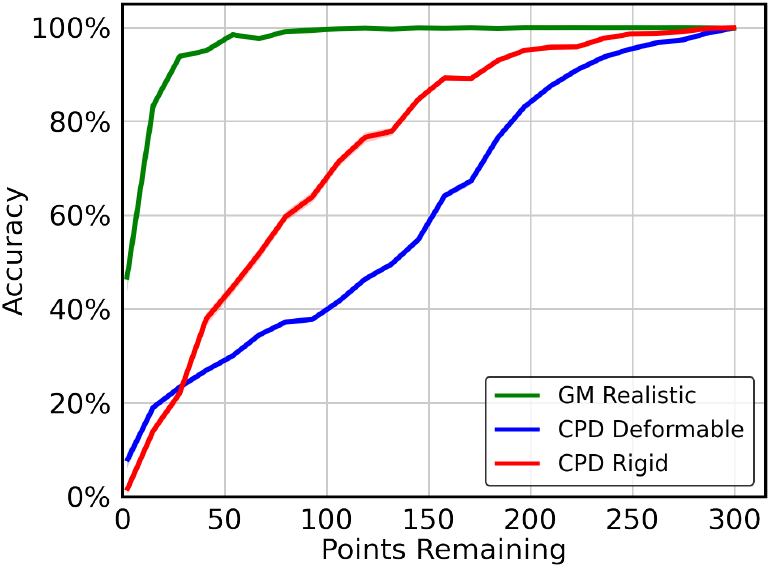
Dropout Performance (40 run average): Alignment accuracy after aligning the OpenWorm atlas to the OpenWorm atlas with random points removed and perturbed as a group in a random direction in x, y, and z each at up to 5 microns.

#### Robustness to Biological Noise

Imaged *C. elegans* individuals do not typically exhibit identical morphology. For example, an individual that has consumed more food may be larger than another individual of the same age. These morphological differences are not isotropic, but their effects on distortion of adjacent neurons present as collective positional shift, so a good identification method must be robust to such local translation. We simulate such “biological noise” as follows.

We randomly generate *N* deformation points **r**_*n*_ within the volume of the neuron point cloud, and a corresponding displacement vector **d**_*n*_ drawn from a 3D Gaussian distribution of standard deviation *σ*, characterizing the “influence radius” of a deformation center, as illustrated in Fig. 5. In other words, each neuron is perturbed according to equation (6), with the only difference that now each **p**_*n*_ and **r**_*n*_ are randomly generated instead of learnable parameters.

This biological noise was applied to our test point cloud according to our realistic noise operation Parameterizing more realistic worm deformation, with *N* = 100 deformation centers, amplitude |**d**_*n*_| = 6.1 and varying influence radius *σ*. We then perturb the larger atlas as a rigid group away from the head neurons by a single vector drawn from a uniform distribution between ±5 microns in each of the *x*, *y*, and *z* directions. Then we use the alignment algorithms to align the unlabeled set to the complete atlas.

We can see that the accuracy of CPD-rigid drops rapidly for biological noise influence radius beyond 15*μm*. This may be because CPD-rigid assumes that all the neurons move as a rigid body that only allows rotation and translation as a whole, which is insufficient to model the realistic biological noise with relative expansion and contraction between neurons. On the other hand, while CPD-deformable performs much better than CPD-rigid, it underperforms GM-realistic across all the tested influence radii of the biological noise.

### Testing alignment methods - Real Data

#### Atlas performance tests

We now test our NeuroPAL-based identification pipeline against previous atlas-based methods, in order to test their comparative performance, and also address the question of how many colors are necessary to achieve acceptable accuracy. In Fig.9–11, we plot the performance of the three tested algorithms on our three tested atlases with a range of numbers of colors. In addition, we include a version of the NeuroPal atlas that has been straightened in a previous way as described by ***Peng et al. (2008)***. In these tests, all atlases were isotropically rescaled so as to occupy *x* ∈ [0*μm*, 800*μm*], and for the midbody tests the atlases were cropped at *x* ∈ [200*μm*, 700*μm*]. Then, alignments were performed using the same parameters as identified in Alignment performance on simulated data.

**Figure 9.**
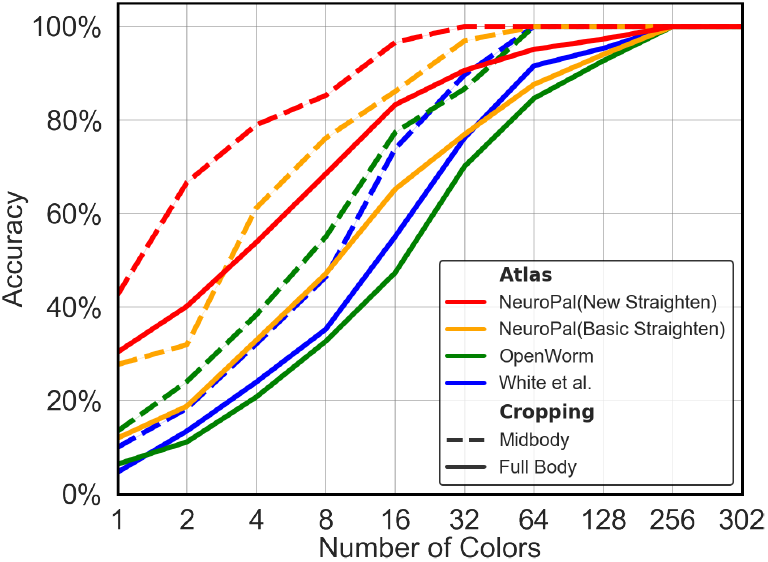
CPD Rigid Alignment (40 run average): Alignment accuracy using CPD Rigid with various atlases and number of colors. Dashed lines denote tests wherein the head and tail of the atlas were cropped out.

**Figure 10.**
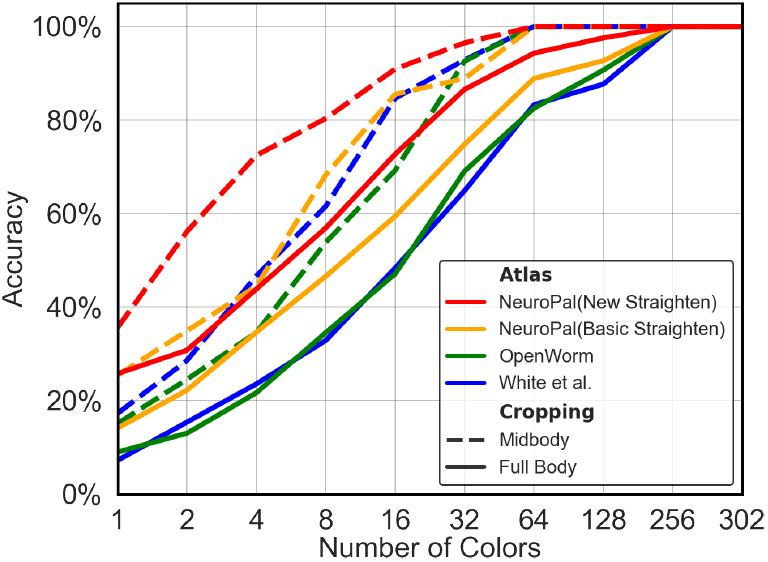
CPD Deformable Alignment (40 run average): Alignment accuracy using CPD Deformable with various atlases and number of colors. Dashed lines denote tests wherein the head and tail of the atlas were cropped out.

**Figure 11.**
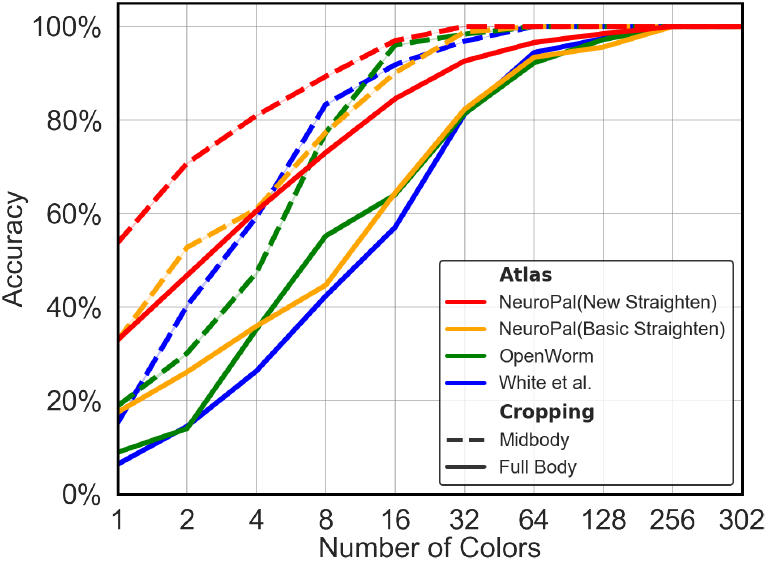
GM Realistic Alignment (40 run average): Alignment accuracy using GM Realistic with various atlases and number of colors. Dashed lines denote tests wherein the head and tail of the atlas were cropped out.

From the results, we have the following observations. Firstly, the NeuroPAL atlas outperforms the OpenWorm and White atlases by a large margin, for all alignment methods, on both full-body and mid-body data. This suggests that our NeuroPAL atlas combined with the circle straightening method enables more accurate identification. Secondly, comparing Fig. 11 and Fig. 9, we see that the GM-realistic method outperforms CPD methods across all colors and atlases and for both full-body and mid-body data, again demonstrating that our alignment method allows more accurate neuron identification. Thirdly, as the number of colors increases, the performance of all atlases increases, emphasizing the important role that NeuroPAL can play in enabling more accurate automated neuron identification for atlas creation.

## Discussion

The main contributions of this paper are

1. a pipeline for constructing a 3D *C. elegans* atlas based on optically imaged neuron data,
2. an alignment method for identification of unlabeled *C. elegans* neurons using such an atlas, and
3. to the best of our knowledge, the most complete *C. elegans* 3D positional neuron atlas, encapsulating positional variability derived from the whole bodies of 7 animals.

We have presented tests suggesting that both our alignment algorithm and our pipeline-produced 3D atlas achieve higher identification accuracy than existing alternatives.

Many groups around the world are in the process of producing better imaging datasets so as to enable more promising investigation of myriad aspects of *C. elegans*, from neuronal activity to gene expression and cell-fate. We hope that, by delivering higher cell-type identification confidence, our atlas and others created using this method will help maximize the scientific value enabled by such functional imaging work.

## Acknowledgments

The authors wish to thank Liam Paninski for helpful comments. E.Y. was funded in part by the NIH (5T32DK7328-37, 5T32DK007328-35, 5T32MH015174-38, and 5T32MH015174-37). E.S.B. was supported by HHMI, Lisa Yang, John Doerr, and NSF Grant 1734870. M.T. was supported by the Foundational Questions Institute, the Rothberg Family Fund for Cognitive Science, and IAIFI through NSF grant PHY-2019786.

## Appendix 1 Hyperparameter settings

Here, we provide the hyperparameters chosen for the GM-realistic and CPD-deformable algorithms, which have been optimized on the experiments in Robustness to Cropping and applied to all experiments. For GM-Realistic, the exponent *γ* in Eq. (4) is set to −5. We train for *T* = 5000 epochs with learning rate lr = 6.31 × 10−^4^ which is reduced by a factor of 0.3 if the best validation loss does not improve for 100 consecutive epochs (30 epochs for the experiment in Robustness to Cropping). To prevent numerical errors from division by nearzero during optimization when 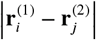 is small (so that 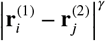 is large, since *γ* < 0), we clamp its value by replacing 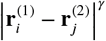 with 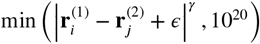 with *ε* = 10^−10^. Moreover, we add a small regularization term that encourages the unlabeled point cloud to keep its original shape, by regularizing Eq. (4) as follows:

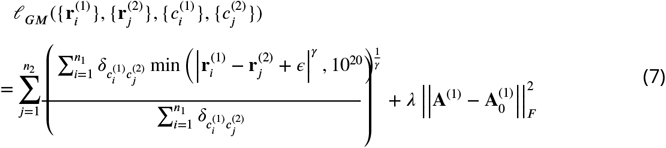

Here **A**^(1)^ is the matrix containing pairwise distances between each neuron pair in **r**^(1)^, and 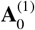, is **A**^(1)^ at the start of training. ||·||_*F*_ denotes Frobenius norm. For *λ*, we use an annealing procedure of 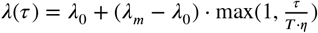 where *τ* is the epoch number, and *λ*_0_, *λ_m_*, *η* are hyperparameters. In the experiment at Testing alignment methods - Real Data, we use *N* = 200 deformation centers, with *σ* = 0.2 and initial amplitude |**d**_0_| of **d** being 0.015. In experiment at Robustness to Cropping, we use *N* = 50 deformation centers, with *σ* = 50 and initial amplitude |**d**_0_| of 0.1, due to different scenario and units. The detailed hyperparameters that we tuned, including search range and the chosen value, is provided in Table 1.

Table 1 below provides the hyperparameter setting. The parameters *α* and *β* are defined in (***Myronenko and Song, 2010***). For other hyperparameters, we use the default setting as in its implementation at https://github.com/siavashk/pycpd (as of Aug 1, 2020).

**Appendix 1 Table 1.**
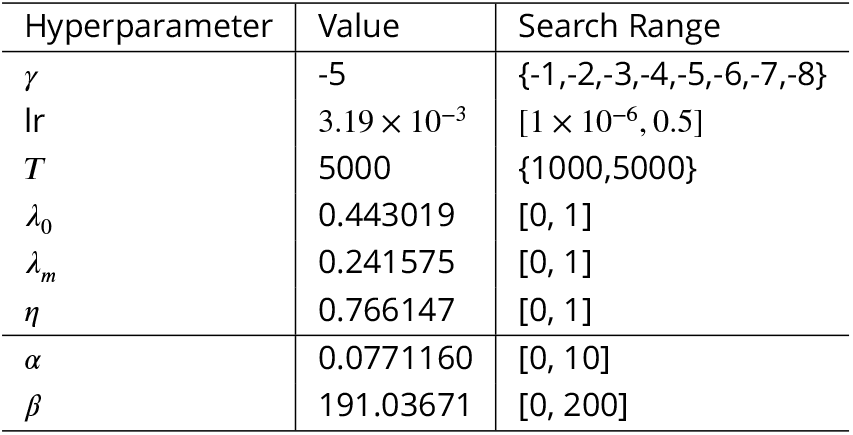
Hyperparameters for the GM-Realistic method (first group) and CPD-Deformable method (*α, β*).

## Appendix 2 The NeuroPAL atlas, tabulated

Here, we provide the neuron coordinates for the NeuroPAL atlas that we generated over the course of this work. The coordinates of these 300 neurons can also be downloaded at https://github.com/bluevex/elegans-atlas. As explained in the text, the two CAN cells are not included in our atlas.

**Appendix 2 Table 1.** Positions of aggregated NeuroPAL atlas, in microns.

| Neuron | x | y | z |
| --- | --- | --- | --- |
| ADAL | 103.47 | -2.24 | -6.33 |
| ADAR | 98.37 | 0.96 | 8.26 |
| ADEL | 93.11 | -4.22 | -6.33 |
| ADER | 68.78 | 3.62 | 6.57 |
| ADFL | 56.05 | 5.07 | -4.98 |
| ADFR | 62.05 | 12.27 | 5.34 |
| ADLL | 65.15 | 11.51 | -4.25 |
| ADLR | 72.58 | 11.16 | 4.29 |
| AFDL | 63.04 | -2.90 | -0.53 |
| AFDR | 57.73 | 8.74 | 5.26 |
| AIAL | 62.96 | 5.22 | -1.75 |
| AIAR | 83.26 | -5.69 | 3.07 |
| AIBL | 66.37 | -0.18 | -4.56 |
| AIBR | 63.35 | 2.45 | 5.18 |
| AIML | 93.83 | -13.30 | -0.82 |
| AIMR | 81.69 | -6.74 | 2.80 |
| AINL | 73.97 | 5.99 | -7.66 |
| AINR | 77.45 | 8.77 | 5.85 |
| AIYL | 75.40 | -4.89 | -3.26 |
| AIYR | 66.75 | -2.42 | 1.85 |
| AIZL | 78.82 | -1.47 | -5.09 |
| AIZR | 76.56 | 2.36 | 5.26 |
| ALA | 54.88 | 17.96 | -1.01 |
| ALML | 338.65 | 11.69 | -11.76 |
| ALMR | 314.65 | 19.82 | 10.80 |
| ALNL | 777.53 | 15.69 | -4.42 |
| ALNR | 787.61 | 14.13 | 2.48 |
| AQR | 91.22 | -0.76 | 6.86 |
| AS1 | 133.21 | -13.15 | 0.74 |
| AS10 | 653.28 | -7.83 | -0.76 |
| AS11 | 721.41 | -5.65 | -1.56 |
| AS2 | 183.96 | -13.03 | -0.37 |
| AS3 | 243.82 | -13.01 | 1.36 |
| AS4 | 315.51 | -15.79 | 0.51 |
| AS5 | 374.68 | -16.37 | 0.11 |
| AS6 | 416.98 | -13.60 | 0.36 |
| AS7 | 479.98 | -12.75 | 1.69 |
| AS8 | 534.79 | -13.08 | -0.11 |
| AS9 | 597.87 | -11.32 | -0.09 |
| ASEL | 67.82 | 5.40 | -7.38 |
| ASER | 68.55 | 10.74 | 6.22 |
| ASGL | 67.93 | 7.44 | -4.75 |
| ASGR | 65.38 | 11.39 | 4.05 |
| ASHL | 63.23 | 2.92 | -6.34 |
| ASHR | 62.97 | 7.72 | 7.46 |
| ASIL | 72.45 | 4.94 | -4.60 |
| ASIR | 63.79 | 14.61 | 1.90 |
| ASJL | 75.85 | -2.09 | -3.74 |
| ASJR | 91.31 | -6.30 | -3.44 |
| ASKL | 54.85 | 3.47 | -4.11 |
| ASKR | 63.18 | 8.14 | 3.91 |
| AUAL | 66.02 | -1.81 | -6.27 |
| AUAR | 68.32 | 0.98 | 4.74 |
| AVAL | 52.34 | 6.23 | -6.66 |
| AVAR | 54.59 | 2.88 | 3.54 |
| AVBL | 71.54 | 5.06 | -3.45 |
| AVBR | 72.77 | 11.02 | 8.07 |
| AVDL | 76.23 | -0.36 | -6.66 |
| AVDR | 77.25 | 5.64 | 6.09 |
| AVEL | 62.61 | 2.09 | -5.27 |
| AVER | 67.33 | 3.84 | 5.10 |
| AVFL | 119.50 | -9.32 | -1.68 |
| AVFR | 117.42 | -10.84 | 0.92 |
| AVG | 120.73 | -10.38 | 1.77 |
| AVHL | 66.15 | 10.31 | -6.76 |
| AVHR | 67.38 | 13.54 | 5.22 |
| AVJL | 70.09 | 7.02 | -6.18 |
| AVJR | 71.74 | 12.39 | 3.80 |
| AVKL | 97.68 | -7.51 | -3.19 |
| AVKR | 98.66 | -8.58 | 3.22 |
| AVL | 60.23 | -1.09 | 2.98 |
| AVM | 287.96 | -1.15 | 10.07 |
| AWAL | 69.04 | 4.23 | -5.48 |
| AWAR | 70.50 | 3.72 | 5.88 |
| AWBL | 60.44 | 7.94 | -6.27 |
| AWBR | 63.66 | 12.65 | 5.84 |
| AWCL | 62.15 | -0.45 | -4.84 |
| AWCR | 67.11 | -0.31 | 4.61 |
| BAGL | 40.21 | 5.32 | -4.85 |
| BAGR | 42.15 | 4.00 | 5.96 |
| BDUL | 188.68 | 7.58 | -10.04 |
| BDUR | 187.79 | 11.41 | 7.80 |
| CEPDL | 52.69 | 15.61 | -4.65 |
| CEPDR | 59.47 | 16.56 | 3.44 |
| CEPVL | 50.12 | 0.83 | -4.10 |
| CEPVR | 42.17 | -0.14 | 2.44 |
| DA1 | 132.48 | -12.43 | -0.36 |
| DA2 | 192.83 | -14.26 | -0.02 |
| DA3 | 268.10 | -14.80 | -0.37 |
| DA4 | 348.45 | -17.81 | -0.58 |
| DA5 | 460.74 | -14.05 | 1.55 |
| DA6 | 545.84 | -13.75 | -0.03 |
| DA7 | 655.63 | -9.00 | -0.73 |
| DA8 | 742.54 | -1.67 | -1.20 |
| DA9 | 742.77 | -1.10 | 0.67 |
| DB1 | 128.74 | -10.25 | 1.48 |
| DB2 | 108.68 | -11.66 | -0.23 |
| DB3 | 181.21 | -14.37 | -0.76 |
| DB4 | 306.22 | -16.12 | -0.36 |
| DB5 | 413.59 | -14.06 | 0.86 |
| DB6 | 532.42 | -14.45 | -0.14 |
| DB7 | 648.91 | -9.61 | -0.77 |
| DD1 | 128.02 | -12.47 | 0.77 |
| DD2 | 226.98 | -13.41 | 1.25 |
| DD3 | 352.61 | -15.46 | 1.37 |
| DD4 | 476.87 | -14.14 | 2.41 |
| DD5 | 590.72 | -11.77 | -0.19 |
| DD6 | 741.67 | -1.61 | -0.23 |
| DVA | 765.38 | 13.46 | 0.45 |
| DVB | 766.31 | 15.01 | -2.16 |
| DVC | 769.17 | 14.37 | -1.98 |
| FLPL | 83.45 | 2.06 | -6.66 |
| FLPR | 81.42 | -1.36 | 5.07 |
| HSNL | 462.61 | -1.76 | -9.73 |
| HSNR | 452.19 | -3.30 | 10.45 |
| I1L | 4.18 | 3.62 | -4.02 |
| I1R | 0.00 | 4.19 | 1.70 |
| I2L | 20.43 | 1.17 | -4.46 |
| I2R | 12.08 | 5.00 | 3.77 |
| I3 | 15.87 | 17.67 | -0.34 |
| I4 | 74.25 | 17.99 | -3.35 |
| I5 | 83.43 | -4.95 | 1.61 |
| I6 | 75.02 | 14.23 | -2.65 |
| IL1DL | 36.20 | 13.98 | 1.18 |
| IL1DR | 31.78 | 12.25 | -0.03 |
| IL1L | 32.30 | 5.06 | -5.29 |
| IL1R | 30.90 | 5.83 | 5.26 |
| IL1VL | 29.02 | -1.00 | -2.67 |
| IL1VR | 43.69 | -3.04 | 0.90 |
| IL2DL | 23.49 | 14.65 | -3.32 |
| IL2DR | 27.04 | 16.15 | 3.36 |
| IL2L | 27.76 | 1.77 | -4.62 |
| IL2R | 30.67 | 9.78 | 5.29 |
| IL2VL | 37.93 | -1.66 | -5.18 |
| IL2VR | 31.50 | 1.42 | 4.48 |
| LUAL | 773.45 | 8.42 | -3.59 |
| LUAR | 772.58 | 6.84 | 2.74 |
| M1 | 89.07 | 14.28 | 6.10 |
| M2L | 89.94 | -2.57 | 0.54 |
| M2R | 79.51 | 8.47 | 4.97 |
| M3L | 25.64 | 3.93 | -4.16 |
| M3R | 34.73 | 2.38 | -1.84 |
| M4 | 24.33 | 10.47 | -0.28 |
| M5 | 98.31 | 9.32 | 0.59 |
| MCL | 5.91 | -0.22 | -2.93 |
| MCR | 22.79 | 2.06 | 2.54 |
| MI | 13.14 | 15.53 | 0.57 |
| NSML | 23.89 | 0.34 | -4.65 |
| NSMR | 16.91 | 0.45 | 2.92 |
| OLLL | 33.24 | 10.30 | -4.20 |
| OLLR | 34.03 | 10.06 | 5.02 |
| OLQDL | 37.33 | 14.50 | -1.40 |
| OLQDR | 38.51 | 14.24 | 2.52 |
| OLQVL | 43.31 | 3.52 | -5.31 |
| OLQVR | 45.70 | -1.79 | 2.03 |
| PDA | 752.57 | -1.23 | -1.14 |
| PDB | 748.23 | -2.11 | -0.01 |
| PDEL | 561.02 | 12.04 | -7.47 |
| PDER | 554.68 | 11.39 | 10.37 |
| PHAL | 767.63 | 7.67 | -4.06 |
| PHAR | 772.70 | 2.41 | 2.05 |
| PHBL | 770.31 | 6.22 | -5.05 |
| PHBR | 771.33 | 4.96 | 3.52 |
| PHCL | 784.57 | 6.50 | -1.26 |
| PHCR | 780.48 | 4.19 | 2.06 |
| PLML | 800.00 | 6.19 | -1.50 |
| PLMR | 423.80 | 7.15 | -0.87 |
| PLNL | 766.47 | 2.71 | -5.51 |
| PLNR | 769.47 | 2.61 | 2.91 |
| PQR | 775.68 | 5.64 | -5.11 |
| PVCL | 777.68 | 9.20 | -4.85 |
| PVCR | 775.69 | 5.24 | 3.91 |
| PVDL | 555.26 | 12.55 | -8.04 |
| PVDR | 539.69 | 11.50 | 6.94 |
| PVM | 555.38 | 9.87 | -7.67 |
| PVNL | 795.88 | 9.77 | -3.60 |
| PVNR | 799.05 | 10.60 | 0.68 |
| PVPL | 732.00 | -3.14 | -1.67 |
| PVPR | 739.86 | -2.67 | -0.63 |
| PVQL | 763.11 | 6.80 | -4.29 |
| PVQR | 769.09 | 5.76 | 3.34 |
| PVR | 787.21 | 7.44 | 0.68 |
| PVT | 731.94 | -4.35 | 0.90 |
| PVWL | 781.33 | 6.89 | -4.96 |
| PVWR | 781.18 | 8.55 | 2.50 |
| RIAL | 51.78 | 11.31 | -4.82 |
| RIAR | 55.40 | 8.43 | 5.75 |
| RIBL | 79.90 | 0.06 | -5.46 |
| RIBR | 69.89 | -0.85 | 0.62 |
| RICL | 83.39 | -1.96 | -5.07 |
| RICR | 74.89 | -3.08 | 4.49 |
| RID | 70.54 | 15.52 | 3.64 |
| RIFL | 100.71 | -11.45 | 0.82 |
| RIFR | 133.94 | -12.21 | 1.22 |
| RIGL | 127.23 | -10.24 | -0.28 |
| RIGR | 119.95 | -9.08 | 0.51 |
| RIH | 49.77 | 2.12 | 4.69 |
| RIML | 70.30 | -0.94 | -3.80 |
| RIMR | 67.28 | -6.29 | 2.75 |
| RIPL | 39.60 | 5.26 | -6.60 |
| RIPR | 39.80 | 8.56 | 4.73 |
| RIR | 51.30 | -4.84 | 0.57 |
| RIS | 91.90 | -10.80 | -2.11 |
| RIVL | 73.60 | 6.93 | -5.48 |
| RIVR | 56.64 | 14.28 | 3.74 |
| RMDDL | 63.00 | -4.22 | -2.98 |
| RMDDR | 69.88 | -2.04 | 2.38 |
| RMDL | 55.61 | 4.64 | -5.99 |
| RMDR | 56.37 | 1.28 | 4.90 |
| RMDVL | 50.29 | 6.26 | -5.89 |
| RMDVR | 52.70 | 8.43 | 5.60 |
| RMED | 40.31 | 16.39 | -0.10 |
| RMEL | 42.65 | 6.14 | -4.44 |
| RMER | 41.20 | 9.41 | 6.22 |
| RMEV | 52.47 | -2.21 | 0.00 |
| RMFL | 64.83 | -6.73 | -0.81 |
| RMFR | 67.73 | -3.71 | 0.55 |
| RMGL | 96.44 | -2.51 | -6.24 |
| RMGR | 99.36 | -0.88 | 5.55 |
| RMHL | 69.53 | -0.16 | 1.86 |
| RMHR | 65.18 | -7.36 | 1.37 |
| SAADL | 59.07 | -2.51 | -1.13 |
| SAADR | 52.59 | -2.67 | 0.67 |
| SAAVL | 52.21 | 10.07 | -5.55 |
| SAAVR | 52.46 | 8.82 | 5.76 |
| SABD | 129.68 | -9.85 | 1.95 |
| SABVL | 110.46 | -7.62 | 1.01 |
| SABVR | 94.41 | -7.88 | 2.51 |
| SDQL | 558.50 | 8.92 | -7.39 |
| SDQR | 243.25 | 19.83 | 9.80 |
| SIADL | 52.88 | 5.99 | -4.74 |
| SIADR | 94.26 | -6.92 | 0.07 |
| SIAVL | 64.90 | -0.57 | 1.00 |
| SIAVR | 74.03 | -3.93 | 2.28 |
| SIBDL | 59.92 | -0.62 | -3.41 |
| SIBDR | 52.27 | 4.06 | 2.84 |
| SIBVL | 70.85 | -3.99 | -2.89 |
| SIBVR | 59.55 | -4.35 | 3.26 |
| SMBDL | 100.41 | -8.18 | 1.69 |
| SMBDR | 69.95 | -4.52 | 2.10 |
| SMBVL | 95.74 | -5.46 | -0.46 |
| SMBVR | 74.54 | -6.54 | -0.34 |
| SMDDL | 59.99 | -2.64 | -3.87 |
| SMDDR | 58.33 | -2.51 | 1.61 |
| SMDVL | 53.28 | 14.26 | -4.33 |
| SMDVR | 53.12 | 14.21 | 4.62 |
| URADL | 28.69 | 15.57 | -2.61 |
| URADR | 31.47 | 13.46 | 3.99 |
| URAVL | 39.31 | 1.52 | -3.45 |
| URAVR | 35.49 | -0.83 | 3.70 |
| URBL | 36.05 | 4.63 | -6.52 |
| URBR | 37.07 | 9.15 | 7.78 |
| URXL | 52.67 | 14.08 | -2.00 |
| URXR | 52.94 | 15.23 | 1.90 |
| URYDL | 34.12 | 14.01 | -3.37 |
| URYDR | 36.45 | 14.87 | 4.10 |
| URYVL | 35.44 | 4.01 | -5.63 |
| URYVR | 42.26 | 7.51 | 4.15 |
| VA1 | 109.56 | -10.77 | -1.31 |
| VA10 | 631.93 | -10.73 | -0.85 |
| VA11 | 689.04 | -7.90 | -1.30 |
| VA12 | 739.99 | -1.88 | -1.51 |
| VA2 | 156.13 | -13.88 | 0.82 |
| VA3 | 211.73 | -14.23 | -0.14 |
| VA4 | 276.38 | -15.94 | 0.64 |
| VA5 | 338.48 | -16.82 | 1.40 |
| VA6 | 408.54 | -13.61 | 2.05 |
| VA7 | 459.29 | -14.80 | 2.91 |
| VA8 | 505.45 | -14.17 | 0.24 |
| VA9 | 571.62 | -13.06 | -0.52 |
| VB1 | 107.58 | -10.59 | 2.36 |
| VB10 | 580.62 | -13.56 | -0.69 |
| VB11 | 639.06 | -9.61 | -0.65 |
| VB2 | 104.95 | -9.32 | -0.25 |
| VB3 | 163.81 | -13.59 | 0.26 |
| VB4 | 219.49 | -14.67 | -0.10 |
| VB5 | 289.05 | -16.23 | 0.62 |
| VB6 | 343.80 | -16.63 | 1.44 |
| VB7 | 406.06 | -14.38 | 1.09 |
| VB8 | 467.44 | -14.92 | 2.96 |
| VB9 | 524.53 | -14.33 | -0.04 |
| VC1 | 226.67 | -13.11 | -0.14 |
| VC2 | 295.79 | -17.33 | 0.49 |
| VC3 | 384.20 | -16.46 | 1.39 |
| VC4 | 442.02 | -11.40 | 0.18 |
| VC5 | 455.70 | -8.41 | 0.57 |
| VC6 | 529.95 | -12.04 | 0.58 |
| VD1 | 119.85 | -10.48 | 0.66 |
| VD10 | 601.82 | -11.12 | -0.04 |
| VD11 | 659.80 | -7.43 | -0.74 |
| VD12 | 727.76 | -4.28 | -1.44 |
| VD13 | 744.62 | -1.44 | 0.90 |
| VD2 | 132.21 | -11.82 | 0.21 |
| VD3 | 209.27 | -13.48 | -0.45 |
| VD4 | 265.48 | -14.48 | 1.17 |
| VD5 | 321.24 | -16.36 | 0.56 |
| VD6 | 384.51 | -15.71 | 0.95 |
| VD7 | 431.96 | -14.12 | -1.04 |
| VD8 | 482.46 | -13.10 | 2.57 |
| VD9 | 547.98 | -13.36 | -0.01 |

